# Quantum-assisted genomic prediction of blast resistance in rice (Oryza sativa L.) using SNP and haplotype representations

**DOI:** 10.64898/2026.11.18.767717

**Authors:** Roberto Jose Guerrero, Jason Alejandro Castano, Félicien Akohoue, Manuel Narvaez, Gloria Mosquera, Angela Joseph Fernando, Michael Gomez Selvaraj

## Abstract

Rice blast caused by *Magnaporthe oryzae* threatens rice production, and genomic prediction may accelerate identification of resistant germplasm. We evaluated classical machine-learning and simulated quantum-kernel models for binary and three-class prediction of leaf blast (BL) and panicle blast (PB) resistance in 204 rice accessions with 2,120 repeated phenotypic observations. Models used 328,597 single-nucleotide polymorphisms (SNPs), 104,068 phased haplotypes, or 49,451 unphased haplotypes. Binary prediction outperformed three-class prediction. The highest binary macro-F1 scores were 0.87 for BL and 0.73 for PB. Phased and unphased haplotypes reduced predictor dimensionality by 68.3% and 85.0%, respectively, while retaining broadly comparable performance. Quantum support vector classification achieved competitive results in selected binary tasks but did not consistently exceed the strongest classical models. These findings establish a benchmark for SNP- and haplotype-based prediction of rice blast resistance and for evaluation of quantum machine learning in crop genomics.

**Core Ideas:**

- Genomic prediction of rice blast resistance was evaluated using 204 rice accessions and repeated field observations.
- Binary genomic classification of phenotypic resistance classes outperformed three-class classification, with leaf blast more predictable than panicle blast.
- .• Phased and unphased haplotypes reduced predictor dimensionality by 68.3% and 85.0%, respectively, with broadly comparable predictive performance.
- Simulated quantum support vector classification was competitive in selected binary tasks but did not consistently outperform leading classical models.

**Plain Language Summary:** Rice blast can affect leaves and panicles which reduce rice production. We used genome-wide DNA markers from 204 rice accessions to predict resistance at both growth stages. Models using individual DNA markers and haplotypes gave similar results, while haplotypes used fewer predictors. Leaf-blast resistance was more predictable than panicle-blast resistance. A simulated quantum classifier was competitive in some binary prediction tasks, but it did not consistently exceed the best conventional machine-learning models. These results support genomic prediction as a complement to field disease screening in rice breeding.

## 1 INTR**ODUCTION**

Rice blast, caused by the fungal pathogen Magnaporthe oryzae, remains one of the most destructive diseases of rice (Oryza sativa L.), posing a major threat to sustainable rice production and global food security. The disease affects multiple plant organs and developmental stages, with leaf blast (BL) primarily occurring during vegetative growth and panicle blast (PB) affecting reproductive tissues, potentially causing substantial grain yield losses (Asibi et al., 2019; Skamnioti & Gurr, 2009). Although BL and PB resistance may share components of their genetic architecture, they represent distinct but interconnected breeding targets. Differences in developmental stage, tissue susceptibility, pathogen populations, and environmental conditions influence disease expression, meaning that resistance to leaf blast does not necessarily confer equivalent resistance to panicle blast (Cheng et al., 2024; Skamnioti & Gurr, 2009; Yu et al., 2022). Developing cultivars with stable and durable resistance therefore requires genomic approaches capable of characterizing and predicting disease responses across both vegetative and reproductive stages.

Recent advances in rice genomics have provided important insights into the genetic architecture of blast resistance and opportunities to improve genomic prediction. Akohoue et al. (2026) investigated BL and PB resistance in the single-seed-descent SSD Tropics and 3K rice populations using genome-wide association studies (GWAS) and GWAS-informed weighted genomic best linear unbiased prediction (GBLUP). The authors reported moderate genetic correlations between BL and PB severity (0.43–0.44), suggesting partially shared but distinct genetic control. Significant marker–trait associations were identified near major blast resistance loci, including the Pi2/Pi9 cluster and Pi33. Incorporating GWAS-derived marker weights improved genomic predictive ability by 4–37% relative to conventional unweighted GBLUP, demonstrating the potential value of integrating biologically informative genomic features into resistance prediction.

Complementing these population-level analyses, Martin et al. (2026) generated chromosome-scale genome assemblies of two genetically divergent rice cultivars, CT21375-F4-4 (CT21) and Moroberekan, and characterized allelic and structural variation at blast resistance loci. Despite exhibiting comparable resistance under multiyear field conditions, the cultivars displayed distinct resistance-associated genomic architectures. CT21 contained several exact or near-exact matches to characterized blast resistance genes, whereas Moroberekan predominantly harbored divergent homologs. Structural variation was also identified, including a tandem duplication within the Pish-associated nucleotide-binding leucine-rich repeat (NLR) gene cluster in CT21. These findings demonstrate that similar resistance phenotypes can arise from different allelic and structural configurations and highlight the value of high-quality reference assemblies for investigating genomic diversity and resistance-associated haplotypes (Martin et al., 2026).

Together, these studies emphasize the importance of moving beyond individual marker associations toward genomic representations that better capture the organization and diversity of resistance-associated regions. Single-nucleotide polymorphisms (SNPs), commonly encoded as allele dosages, provide high-resolution information about genomic variation and are widely used in genomic prediction. However, high-density SNP datasets frequently contain extensive linkage disequilibrium (LD), correlated markers, and potentially redundant information. Haplotype-based representations offer a complementary strategy by capturing combinations of alleles inherited together within local genomic regions. Such representations may preserve allelic context, reduce predictor dimensionality, and improve the biological interpretation of genomic variation.

Nevertheless, their predictive benefits depend on haplotype-block definition, population structure, trait architecture, and validation design; fewer predictor variables do not necessarily indicate independent genomic regions or improved prediction accuracy (Bhat et al., 2021; MacNish et al., 2026). Systematic comparisons with SNP-based models are therefore essential.

Machine-learning approaches provide flexible computational frameworks for evaluating alternative genomic representations. Algorithms such as gradient-boosted decision trees, random forests (RF), support vector machines (SVM), and multilayer perceptrons can capture nonlinear relationships between genomic predictors and disease responses (Azodi et al., 2019; Grinberg et al., 2020). These approaches complement conventional statistical genomic prediction methods and can accommodate complex, high-dimensional datasets. However, improved prediction from nonlinear models does not necessarily establish biological epistasis or causal genetic interactions. Predictive associations must therefore be distinguished from experimentally validated resistance mechanisms (Ubbens et al., 2021; Upstill-Goddard et al., 2013). Moreover, model performance depends on validation design, class prevalence, and the information retained after preprocessing, emphasizing the importance of rigorous benchmarking (Tong & Nikoloski, 2021).

Alongside classical machine learning, quantum machine learning (QML) has emerged as an exploratory computational framework for investigating complex data representations. Quantum feature-map methods encode classical input variables into quantum states and evaluate similarities between observations in quantum feature spaces. Havlíček et al. (2019) demonstrated supervised learning using quantum-enhanced feature spaces, establishing an important conceptual foundation for quantum kernel methods, including the quantum support vector classifier (QSVC). Variational quantum algorithms provide a complementary approach in which parameterized quantum circuits are optimized through hybrid quantum–classical procedures (Cerezo et al., 2021). These developments motivate investigations into whether quantum representations can provide useful alternatives or complements to established classical algorithms in biologically relevant prediction tasks.

Nevertheless, applying QML to genome-scale datasets presents substantial computational and methodological challenges. Modern genomic datasets may contain hundreds of thousands or millions of variants, whereas practical quantum implementations remain constrained by available qubits, circuit depth, noise, and computational resources. Direct encoding of complete genome-wide SNP or haplotype matrices is therefore generally impractical with current quantum resources.

Feature selection, dimensionality reduction, and biologically informed genomic aggregation provide potential strategies for connecting high-dimensional genomic information with quantum computation. However, these transformations may remove informative genetic variation, and their effects must be distinguished from those of the quantum classifier itself. Consequently, quantum-kernel performance obtained after feature selection and principal-component reduction should be interpreted as the outcome of the complete analytical pipeline rather than evidence of an isolated quantum computational advantage (Havlíček et al., 2019; Tong & Nikoloski, 2021).

Recent exploratory applications of quantum computing to biological datasets further illustrate these methodological opportunities and challenges. Romero et al. (2026a, 2026b) investigated quantum and hybrid quantum–classical approaches for representing complex relationships in single-cell transcriptomic data. In particular, Romero et al. (2026b) proposed a quantum generative framework employing parameterized quantum circuits to represent nonlinear gene–gene and cell– cell relationships. Although these emerging applications do not establish a general quantum advantage in biological data analysis, they demonstrate growing interest in quantum representations of complex biological systems and reinforce the need for rigorous evaluation in crop genomics.

Beyond genomic prediction, quantum and quantum-inspired optimization may provide future opportunities for investigating higher-order genetic interactions underlying complex disease resistance. Multilocus interactions can contribute to phenotypic variation, but their combinatorial complexity makes exhaustive genome-wide searches challenging. Kikuchi and Tanaka (2026) explored higher-order epistasis detection using multifactor dimensionality reduction combined with factorization-machine-based quadratic optimization annealing (FMQA), providing a simulation-based proof of concept for optimization-guided interaction discovery. Although such approaches have not demonstrated quantum computational advantage or experimentally validated higher-order interactions in crop populations, they suggest opportunities to integrate genomic information, classical artificial intelligence, and quantum-assisted optimization. Rice blast resistance provides a relevant biological context for these future investigations; however, the present study focuses specifically on benchmarking classical and simulated quantum machine learning for SNP- and haplotype-based resistance prediction.

An important unresolved question is therefore not simply whether quantum approaches can outperform classical algorithms, but under which genomic representations, phenotypic classification schemes, and computational conditions quantum-kernel methods might provide reproducible predictive benefits. Addressing this question requires careful comparison of classical and quantum models, explicit consideration of dimensionality reduction, and consistent evaluation across biologically distinct disease traits. Leaf and panicle blast are particularly suitable for such comparative analysis because their partially shared genetic architecture and differing disease-expression patterns may influence genomic predictability.

In the present study, we investigated genomic prediction of BL and PB resistance using a rice diversity panel comprising 204 accessions and 2,120 repeated phenotypic observation records. A common CT21-reference-based genomic dataset containing 328,597 SNP predictor variables provided the foundation for evaluating individual SNP and haplotype representations. Phased and unphased haplotype matrices comprising 104,068 and 49,451 predictor variables, respectively, were compared with the SNP representation. These representations were evaluated under binary and three-class classification schemes using classical and simulated quantum machine-learning approaches. Binary classification distinguished resistant and susceptible responses, whereas three-class classification retained resistant, intermediate, and susceptible categories, allowing assessment of the effects of phenotypic classification resolution on model performance.

The specific objectives were to: (i) benchmark classical machine-learning (ML) and simulated quantum machine-learning (QML) approaches for binary and three-class classification of BL and PB resistance; (ii) compare individual SNP, phased-haplotype, and unphased-haplotype representations derived from a common genomic resource; (iii) determine how genomic representation and phenotype classification resolution influence predictive performance; (iv) investigate differences in genomic predictability between BL and PB resistance; and (v) assess the computational characteristics and practical feasibility of simulated QML relative to classical ML under the constraints of high-dimensional genomic data.

We hypothesized that genomic representation and phenotypic classification resolution would influence predictive performance, with haplotype-based encoding potentially preserving informative local allelic combinations while reducing predictor dimensionality. We further hypothesized that simulated quantum-kernel methods could achieve competitive predictive performance using compressed genomic representations, without assuming an inherent quantum advantage.

Collectively, this study establishes a comparative benchmarking framework for SNP-and haplotype-based rice blast resistance prediction and provides a foundation for future investigations of higher-order genetic interactions and quantum-assisted genomic discovery.

## 2 MATERIALS AND METHODS

### 2.1. Plant Materi**als and Rice Diversity Panel**

The study comprised 204 rice accessions from the 3K Rice Genomes Project diversity panel, for which whole-genome sequencing data and field-based phenotypic evaluations of leaf blast (BL) and panicle blast (PB) resistance were available. Disease-resistance evaluations were conducted between 2020 and 2023 at the Santa Rosa rice blast hotspot in Colombia, providing phenotypic observations under natural field disease pressure. The integrated phenotypic dataset comprised 2,120 observation records collected across repeated evaluations. As individual accessions were evaluated multiple times, these records represented repeated phenotypic observations rather than genetically independent samples.

To ensure consistency across genomic prediction approaches, a common set of 328,597 single-nucleotide polymorphism (SNP) predictors were established. Three genomic representations were subsequently considered: individual SNPs (328,597 predictors), phased haplotypes (104,068 predictors), and unphased haplotypes (49,451 predictors). These representations provided a common genomic framework for comparing classical and quantum machine-learning approaches to blast resistance prediction (Table 1).

**Table 1.**
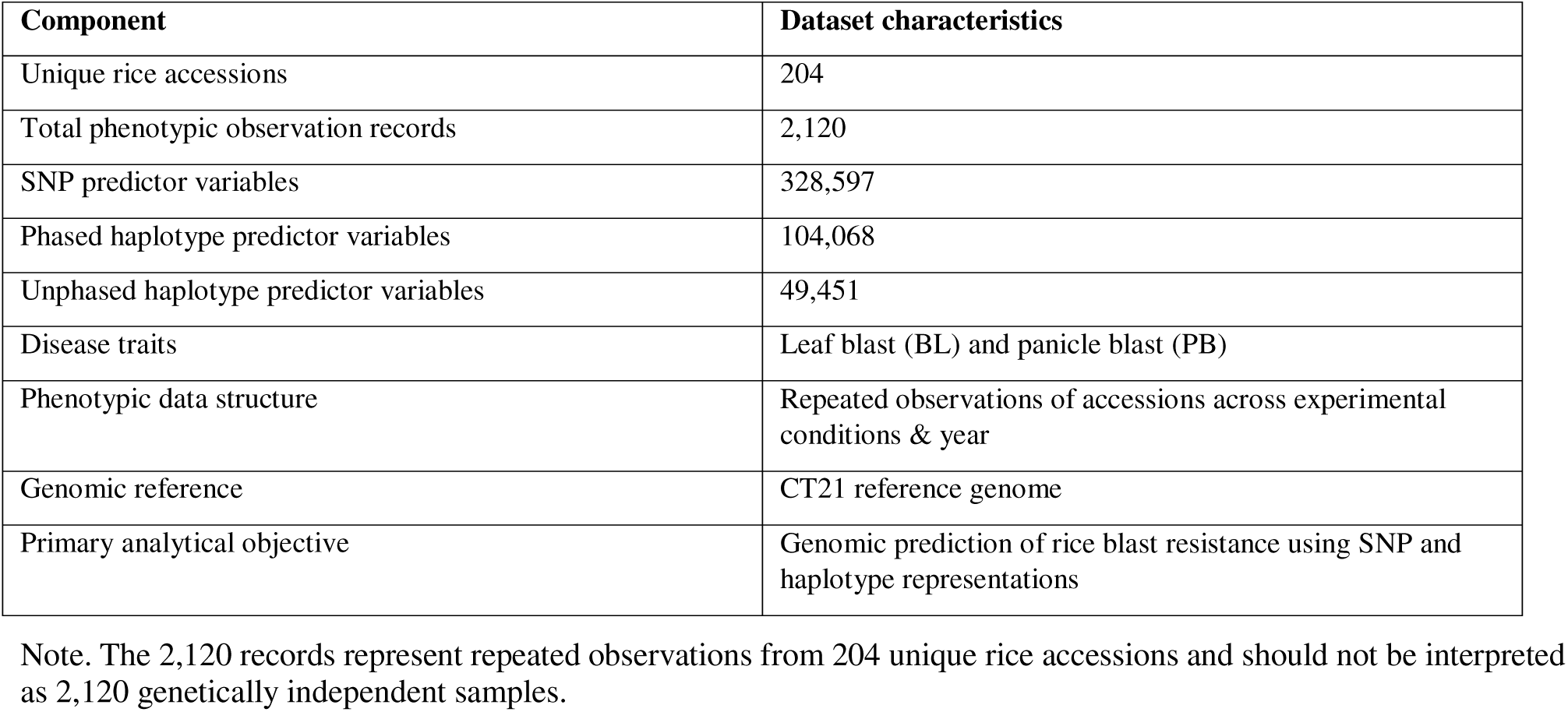
Summary of genomic predictor matrices and phenotypic observations used for rice blast resistance analysis.

| **Component** | **Dataset characteristics** |
|---|---|
| Unique rice accessions | 204 |
| Total phenotypic observation records | 2,120 |
| SNP predictor variables | 328,597 |
| Phased haplotype predictor variables | 104,068 |
| Unphased haplotype predictor variables | 49,451 |
| Disease traits | Leaf blast (BL) and panicle blast (PB) |
| Phenotypic data structure | Repeated observations of accessions across experimental conditions & year |
| Genomic reference | CT21 reference genome |
| Primary analytical objective | Genomic prediction of rice blast resistance using SNP and haplotype representations |

Note. The 2,120 records represent repeated observations from 204 unique rice accessions and should not be interpreted as 2,120 genetically independent samples.

### 2.2. Leaf and Panicle Blast Phenotyping

Leaf blast (BL) and panicle blast (PB) resistance were evaluated separately under field conditions at the Santa Rosa blast hotspot during the 2020–2023 evaluation period. Disease severity was recorded using a nine-point ordinal scale (1–9), with lower scores indicating greater resistance and higher scores indicating increased susceptibility.

To account for the ordinal nature of disease-severity observations and the repeated phenotypic evaluations, a Bayesian ordinal mixed model was used to estimate accession-level disease responses. Maximum a posteriori (MAP) estimates derived from the model were subsequently used to establish the phenotypic targets for genomic prediction. BL and PB were analyzed independently to capture potential differences in their genetic architecture and resistance responses, consistent with previous genomic investigations of rice blast resistance (Akohoue et al., 2026).

### 2.3. Blast Resistance Phenotypic Classification

To assess the influence of phenotypic resolution on genomic prediction performance, MAP-based disease-severity estimates were transformed into two classification schemes: binary and three-class classification. The same classification criteria were applied independently to BL and PB.

For binary classification, accessions were categorized into two resistance groups: resistant (R; scores 1–5) and susceptible (S; scores 6–9). This classification provided a broad distinction between relatively resistant and susceptible disease responses.

For three-class classification, accessions were categorized as resistant (R; scores 1–4), intermediate (I; scores 5–6), and susceptible (S; scores 7–9). This scheme retained an intermediate resistance category, enabling the evaluation of model performance across finer phenotypic distinctions.

Both classification schemes were used to compare classical and quantum machine-learning models across SNP- and haplotype-based genomic representations. This approach enabled an assessment of how phenotype categorization influenced predictive accuracy, class discrimination, and the relative performance of different genomic representations.

### 2.4. Whole-Genome Sequencing and Variant Discovery

Whole-genome sequencing reads for the 204 rice accessions were retrieved from the European Nucleotide Archive (ENA) as paired-end Illumina reads. Sequence data were processed against the CT21 reference genome, restricting variant discovery to the 12 nuclear chromosomes.

Bioinformatic analyses were conducted using a reproducible workflow implemented in Snakemake v8.25.5 (Mölder et al., 2021). Raw sequencing reads were subjected to adapter trimming and quality filtering using Trimmomatic v0.40 (Bolger et al., 2014) in paired-end mode. Quality-filtered reads were aligned to the CT21 reference genome using BWA-MEM v0.7.18 (Li & Durbin, 2009). Sample-specific read-group information was incorporated, and alignments were converted into coordinate-sorted BAM files for downstream processing.

Variant discovery was performed using NGSEPcore v5.0.0 (Tello et al., 2019), following the two-stage variant-calling workflow decribed by Lobatón et al. (2018). First, variants were identified independently for each accession using SingleSampleVariantsDetector, and the resulting calls were combined using MergeVariants to construct a common catalog of variant sites.

Subsequently, all accessions were genotyped at the identified sites using this catalog as a reference, and the resulting genotype calls were consolidated into a multisample VCF using VCFMerge.

SNPs were filtered using a minimum genotype call rate of 0.80 and a minor allele frequency threshold of 0.05, resulting in 3,026,480 high-quality SNPs. This filtered dataset was used independently for selective linkage disequilibrium pruning (SLDP) to construct the individual-SNP predictor matrix and for haplotype-block construction to derive phased and unphased predictor representations.

To obtain the individual-SNP predictor matrix, the 3,026,480-SNP dataset was subjected to SLDP following Akohoue et al. (2026), who adapted the procedure of Zhu et al. (2023). Linkage disequilibrium was measured as the squared allele-frequency correlation (r^2^; Mangin et al., 2012), with the kinship matrix estimated using the VanRaden method. The procedure comprised four steps: (1) identifying GWAS-significant SNPs for each trait and markers within 50 kb in very high LD (r^2^ ≥ 0.95) with them; (2) pruning the significant SNPs and their linked neighbors according to r^2^ and GWAS *P*-values; (3) pruning the remaining markers genome-wide using r^2^ ≥ 0.80; and (4) combining the retained markers while preserving all GWAS-detected SNPs. This procedure generated 328,597 SNP predictors for genomic prediction.

### 2.5. Haplotype Block Construction

Haplotype blocks were constructed from the 3,026,480 filtered SNPs described above. Markers were phased using Beagle 5.x (Browning et al., 2018). Block boundaries were identified chromosome by chromosome using pairwise linkage disequilibrium (*r*^2^) between marker dosages.

Only marker pairs separated by ≤500 kb were evaluated, and pairs with *r*^2^ ≥ 0.70 were considered in high LD. Blocks containing fewer than five SNPs were excluded.

Within each retained block, marker combinations were represented as either phased gametic haplotypes or unphased multi-SNP genotype patterns. Phased haplotype alleles were encoded by copy number (0, 1 or 2), whereas unphased patterns were encoded by absence or presence. Alleles or patterns with a frequency below 0.01 were excluded. This procedure generated 104,068 phased haplotype predictors and 49,451 unphased haplotype-pattern predictors for genomic prediction.

### 2.6 SNP and Haplotype Genomic Representations

Three genomic predictor representations derived from the same 3,026,480-SNP source dataset were evaluated: SLDP-selected individual SNPs, phased haplotypes, and unphased haplotype patterns. The individual-SNP representation comprised 328,597 SLDP-selected markers encoded as alternative-allele dosage, with 0, 1 and 2 representing homozygous reference, heterozygous and homozygous alternative genotypes, respectively. The phased representation comprised 104,068 haplotype-allele predictors that retained the arrangement of alleles on homologous chromosomes.

The unphased representation comprised 49,451 multi-SNP genotype-pattern predictors without phase information.

Relative to the 328,597-predictor SNP matrix, the phased and unphased representations reduced predictor dimensionality by 68.3% and 85.0%, respectively. These percentages describe reductions in the number of predictor columns and not the numbers of independent linkage disequilibrium blocks. All three representations were evaluated for BL and PB resistance prediction using the same classical and quantum machine-learning workflows (Table 1).

### 2.7 Classical and Quantum Machine-Learning Framework

#### 2.7.1 Classical Machine-Learning Class**ifiers**

Classical machine-learning models were evaluated for genomic prediction of leaf blast (BL; Class_MAP_BL) and panicle blast (PB; Class_MAP_NBL) resistance using SNP, phased-haplotype, and unphased-haplotype representations. Prediction targets were derived from Bayesian ordinal mixed-model estimates and evaluated under binary (resistant versus susceptible) and three-class (resistant, intermediate, and susceptible) classification schemes.

Four classifiers were implemented: Extreme Gradient Boosting (XGBoost), Random Forest (RF), Light Gradient Boosting Machine (LightGBM), and Support Vector Machine (SVM) (Breiman, 2001; Chen & Guestrin, 2016; Cortes & Vapnik, 1995; Ke et al., 2017). XGBoost and LightGBM employed gradient-boosted decision trees to capture nonlinear marker–phenotype relationships, RF used bagging-based ensemble learning, and SVM identified class-separating boundaries in high-dimensional feature spaces.

The configured XGBoost model used 100 boosting trees, a maximum depth of five, a learning rate of 0.1, and a random seed of 42. Class-imbalance adjustments and model-specific hyperparameters were applied as appropriate. All classifiers were evaluated separately for BL and PB across both classification schemes and genomic representations, providing classical benchmarks for comparison with quantum machine-learning approaches.

#### 2.7.2 Quantum and Quantum-Inspired Machine-Learning Classifiers

The quantum-learning framework considered four approaches: Quantum Support Vector Classifier (QSVC), Pegasos Quantum Support Vector Classifier (PegasosQSVC), Variational Quantum Classifier (VQC), and Factorization Machine with Quantum Annealing (FMQA), representing quantum-kernel classification, variational learning, and annealing-based optimization (Cerezo et al., 2021; Havlíček et al., 2019; Kikuchi & Tanaka, 2026; Shalev-Shwartz et al., 2007).

QSVC employed a quantum feature map and fidelity kernel for support-vector classification (Havlíček et al., 2019), whereas PegasosQSVC combined quantum-kernel learning with stochastic subgradient optimization (Shalev-Shwartz et al., 2007). VQC used parameterized quantum circuits optimized through a classical algorithm to minimize classification loss (Cerezo et al., 2021). FMQA provided a complementary optimization framework in which a factorization-machine surrogate models feature interactions and an annealing-based optimizer searches for favorable configurations (Kikuchi & Tanaka, 2026). Unlike the other approaches, FMQA is not inherently a supervised classifier and requires a separately defined prediction or feature-selection objective.

To accommodate high-dimensional genomic data, feature selection and dimensionality reduction preceded quantum-circuit processing. Specifically, QSVC employed mutual-information selection of 64 predictors, followed by standardization and reduction to four principal components, encoded using a four-qubit ZZ feature map with two repetitions. Gate-based quantum models were evaluated using classical quantum-circuit simulators. FMQA implementation details and computational backends were documented separately to distinguish simulated, quantum-inspired, and hardware-based computations.

Models were evaluated using consistent resistance-class definitions and comparable validation partitions, with performance assessed through accuracy, balanced accuracy, macro-F1 score, and class-specific metrics.

#### 2.7.3 Feature Selection, Dimensionality Reduction, and Quantum Encoding

Genomic preprocessing was tailored to individual classical and quantum machine-learning algorithms. SNP genotypes were encoded as 0, 1, and 2 for homozygous reference, heterozygous, and homozygous alternative genotypes, respectively. Three-class phenotypes were encoded as resistant (R = 0), intermediate (I = 1), and susceptible (S = 2). Predefined training and testing partitions were established before model-specific preprocessing.

The genomic datasets comprised 328,597 SNPs, 104,068 phased haplotypes, and 49,451 unphased haplotypes. Additional marker-density subsets of approximately 9K, 166K, and 328K SNPs were considered. The 9K dataset was aligned to the Nipponbare reference genome, whereas the 166K and 328K datasets were derived using the CT21375-F4-4 (CT21) reference assembly.

For classical learning, XGBoost retained the complete genomic predictor matrix without additional feature selection or dimensionality reduction. Random Forest, LightGBM, and SVM employed model-specific preprocessing, including scaling or feature reduction where required.

For QSVC, mutual-information ranking selected the 64 highest-scoring genomic predictors using training data only. These features were standardized using a training-fitted StandardScaler and reduced to four principal components through principal component analysis (PCA). The resulting components were encoded using a four-qubit ZZ feature map with two repetitions, followed by quantum fidelity-kernel computation for support-vector classification. PegasosQSVC and VQC used quantum-compatible feature representations with model-specific circuit and optimization configurations. FMQA was considered separately because its factorization-machine surrogate and annealing-based optimization differ from gate-based quantum classification.

All supervised feature selection, scaling, and PCA transformations were fitted exclusively on training partitions and subsequently applied to test data to minimize information leakage. Because classical models could retain substantially more genomic information than quantum models, comparative performance was interpreted in relation to differences in preprocessing, feature dimensionality, and model architecture.

#### 2.7.4 Cross-Validation and Model Performance Evaluation

Classical and quantum machine-learning models were evaluated using a unified framework across genomic representations, disease traits, and classification schemes. Maximum a posteriori (MAP)-derived resistance categories for leaf blast (BL) and panicle blast (PB) were assessed under binary and three-class classification.

Predefined training and testing partitions were used for model evaluation. Given the repeated phenotypic observations of 204 rice accessions, potential overlap of identical accessions between partitions was examined. Accession-grouped cross-validation was identified as the appropriate strategy for evaluating generalization to unseen genotypes. Where available, identical held-out samples were used for classical and quantum models, with supervised preprocessing, including mutual-information feature selection, restricted to training data to minimize information leakage.

Model performance was assessed using accuracy, balanced accuracy, class-specific precision, recall, and F1-score, together with macro-averaged precision, recall, and F1-score. Macro-averaged metrics assigned equal weight to each resistance class, regardless of class frequency. Confusion matrices were used to examine misclassification patterns, while receiver operating characteristic area under the curve (ROC-AUC) and precision–recall area under the curve (PR-AUC) were calculated for binary classification when continuous prediction scores were available.

Binary classification results are presented in Table 2, and three-class results in Supplementary Table S1. Comparisons across genomic representations and computational approaches were interpreted considering validation consistency, feature dimensionality, and model-specific preprocessing.

**Table 2.** Comparative predictive performance of classical and quantum machine learning models using SNP and haplotype representations for rice blast resistance.

| **Genomic representation** | **Model** | **Accuracy (%)** | **Balanced accuracy** | **Macro-F1** |
|---|---|---|---|---|
| SNP (328,597) | XGBoost | 79.6 | 80.0 | 78.5 |
|  | SVM | 80.0 | 80.0 | 79.0 |
|  | Random Forest | 80.0 | 80.0 | 79.0 |
|  | LightGBM | 79.8 | 80.0 | 79.0 |
|  | QSVC | 80.0 | 77.5 | 77.5 |
|  | PegasosQSVC | 61.2 | 50.0 | 38.0 |
|  | VQC | 52.9 | 55.0 | 52.5 |
| Phased haplotypes (104,068) | XGBoost | 80.0 | 80.5 | 79.5 |
|  | SVM | 80.9 | 81.0 | 80.0 |
|  | Random Forest | 80.0 | 80.0 | 79.0 |
|  | LightGBM | 80.2 | 80.5 | 79.0 |
|  | QSVC | 77.5 | 73.5 | 74.0 |
|  | PegasosQSVC | 61.2 | 50.0 | 38.0 |
|  | VQC | 47.7 | 48.0 | 47.0 |
| Unphased haplotypes (49,451) | XGBoost | 79.4 | 79.5 | 78.5 |
|  | SVM | 80.4 | 81.0 | 79.5 |
|  | Random Forest | 80.0 | 80.0 | 79.0 |
|  | LightGBM | 79.8 | 80.0 | 79.0 |
|  | QSVC | 76.5 | 74.0 | 74.0 |
|  | PegasosQSVC | 61.2 | 50.0 | 38.0 |
|  | VQC | 48.1 | 47.0 | 47.0 |

Note. Accuracy, balanced accuracy, and macro-F1 are unweighted means of trait-specific binary test results.

#### 2.7.5 Computational Implementation and Runtime Assessment

Computational analyses were conducted in Python using Scikit-learn, XGBoost, LightGBM, PyCaret, NumPy, and Pandas, with Matplotlib and Seaborn for visualization. Gate-based quantum machine-learning models were implemented using Qiskit Machine Learning. Classical models were executed on conventional computing resources, whereas QSVC, PegasosQSVC, and VQC were evaluated using quantum-circuit simulators. The FMQA workflow was considered separately, with its annealing backend and optimization settings documented according to the computational environment.

Wall-clock times for data loading and model training were recorded by disease trait, classification scheme, genomic representation, and model configuration. Because classical and quantum approaches differed in feature dimensionality, preprocessing, circuit simulation, and optimization requirements, runtime comparisons were interpreted descriptively rather than as evidence of computational speedup or quantum advantage. Software versions, hardware specifications, hyperparameters, quantum-circuit configurations, and optimization settings are detailed in Supplementary Table S4.

## 3 RESULTS

### 3.1. Phenotypic variation and relationship between leaf and panicle blast

Substantial phenotypic variation was observed for leaf blast (BL) and panicle blast (PB) resistance across the rice diversity panel. The dataset comprised 204 unique rice accessions with 2,120 phenotypic observation records obtained from repeated evaluations. Disease responses were assessed using a nine-point severity scale, allowing characterization of resistance variation across vegetative and reproductive developmental stages (Figure 1A–B).

**Figure 1.**
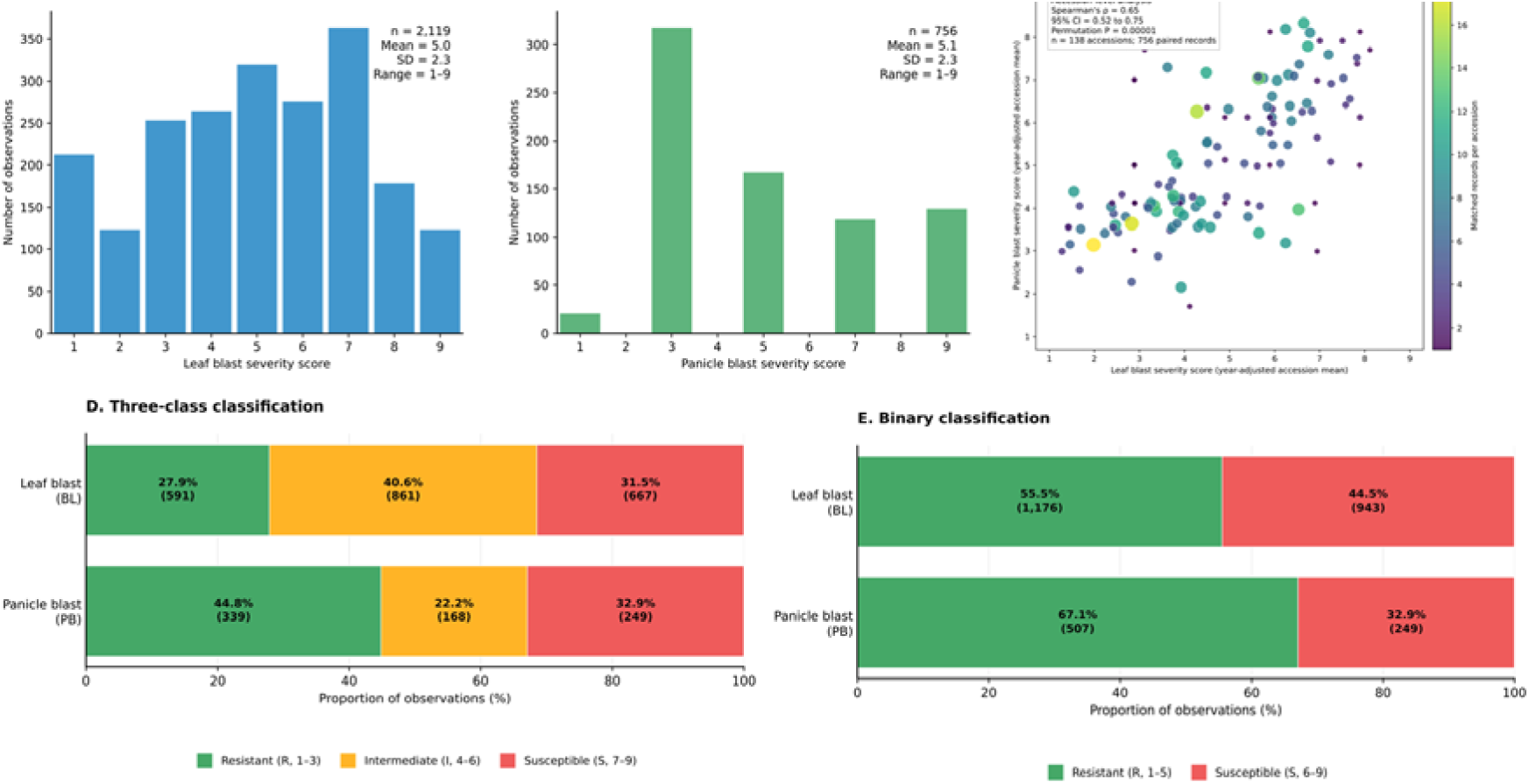
Phenotypic variation and relationship between leaf blast (BL) and panicle blast (PB) resistance in the rice diversity panel. (A–B) Distribution of disease-severity scores for BL and PB, respectively, based on the original nine-point ordinal scale. (C) Year-adjusted accession-level relationship between BL and PB severity (n = 138 accessions; 756 matched records); point color and size indicate the number of matched records per accession. (D) Distribution of resistant (R; scores 1–3), intermediate (I; scores 4–6), and susceptible (S; scores 7–9) phenotypes under the three-class classification scheme. (E) Distribution of resistant (R; scores 1–5) and susceptible (S; scores 6–9) phenotypes under binary classification. Class frequencies are presented separately for BL and PB.

The distribution of BL and PB severity scores was examined to characterize differences in disease response and identify the relative frequencies of resistant, intermediate, and susceptible phenotypes. BL severity ranged from 1 to 9, with a mean ± SD of 5.02 ± 2.29 (n = 2,119), whereas PB severity ranged from 1 to 9, with a mean ± SD of 5.05 ± 2.35 (n = 756). Under the three-class scheme, BL included 591 resistant (27.9%), 861 intermediate (40.6%), and 667 susceptible (31.5%) observations, whereas PB included 339 resistant (44.8%), 168 intermediate (22.2%), and 249 susceptible (32.9%) observations. Thus, BL responses were more concentrated in the intermediate class, while PB showed a higher proportion of resistant observations and a lower proportion of intermediate observations (Figure 1A-B).

To investigate the relationship between resistance at vegetative and reproductive stages, BL and PB severity scores were compared across accessions (Figure 1C). To avoid treating repeated field observations as independent, matched BL-PB records were centered within evaluation year and then averaged for each accession, yielding 138 accession-level estimates from 756 paired records.

Spearman’s rank correlation showed a strong positive phenotypic association between the traits (ρ = 0.65, 95% bootstrap CI = 0.52-0.75; permutation P < 0.00001). This result indicates that accessions with lower BL severity generally also had lower PB severity, while the scatter of accession means identifies accessions with contrasting responses between the two developmental stages.

For genomic prediction, the original nine-point severity scores were transformed into binary and three-class response categories. Under the three-class classification scheme, resistant (R), intermediate (I), and susceptible (S) phenotypes corresponded to scores of 1–4, 5–6, and 7–9, respectively. Under binary classification, scores of 1–5 were categorized as resistant and scores of 6–9 as susceptible. The resulting class distributions were examined separately for BL and PB (Figure 1D–E).

Together, these analyses provide a phenotypic foundation for evaluating the influence of disease trait, resistance classification, and genomic representation on classical and quantum machine-learning prediction performance.

### 3.2. Dataset distribution and partition balance

The supplied ternary class-distribution figures show that the BL partitions were proportionally stable: the training partition contained 414 resistant (27.9%), 602 intermediate (40.6%), and 467 susceptible (31.5%) observations (n = 1,483), while the test partition contained 177 resistant (27.8%), 259 intermediate (40.7%), and 200 susceptible (31.4%) observations (n = 636). The corresponding PB partitions comprised 237 resistant (44.8%), 118 intermediate (22.3%), and 174 susceptible (32.9%) observations in training (n = 529), and 102 resistant (44.9%), 50 intermediate (22.0%), and 75 susceptible (33.0%) observations in testing (n = 227).

Thus, the reported train and test sets retained closely matched ternary class proportions for both traits. BL was enriched for the intermediate category, whereas PB had a larger resistant category and a smaller intermediate category. The binary distribution panels likewise show unequal resistant and susceptible counts, so accuracy is interpreted together with macro-averaged metrics rather than as a standalone indicator of class discrimination (Figure 1).

### 3.3. Comparative Performance of SNP- and Haplotype-Based Genomic Prediction

The predictive performance of classical and quantum machine-learning models was systematically compared using three genomic representations: 328,597 single-nucleotide polymorphisms (SNPs), 104,068 phased haplotype predictors, and 49,451 unphased haplotype predictors. Genomic prediction was evaluated independently for leaf blast (BL) and panicle blast (PB) resistance under binary classification using overall accuracy, balanced accuracy, and macro-F1 score (Table 2). Three-class classification, distinguishing resistant, intermediate, and susceptible phenotypes, was additionally evaluated to assess prediction performance at higher phenotypic resolution (Supplementary Table S1).

Comparative analyses examined whether phased and unphased haplotype representations could maintain or improve predictive performance while substantially reducing genomic predictor dimensionality relative to individual SNPs. Classical and quantum machine-learning models were compared under consistent validation conditions to assess the influence of genomic representation and learning approach on prediction accuracy. Differences between BL and PB were further examined to determine whether predictive performance varied according to disease trait and genomic representation.

For the compiled binary test results, XGBoost achieved BL accuracies of 0.8648, 0.8679, and 0.8648 with SNP, phased-haplotype, and unphased-haplotype predictors, respectively; corresponding PB accuracies were 0.7269, 0.7313, and 0.7225. QSVC achieved BL accuracies of 0.83, 0.84, and 0.82 and PB accuracies of 0.77, 0.71, and 0.71 across the same representations. Across the reported classical model-representation combinations, the highest binary BL accuracy was 0.8726 for SVM with phased haplotypes, and the highest PB accuracy was 0.7445 for the same combination. QSVC achieved a higher PB accuracy of 0.77 for the SNP representation, with a macro-F1 score of 0.73 (Table 2).

Three-class prediction was more difficult, especially for PB. For XGBoost, macro-F1 was 0.69, 0.69, and 0.68 for BL using SNP, phased-haplotype, and unphased-haplotype representations, whereas PB macro-F1 was 0.49, 0.48, and 0.48, respectively. QSVC macro-F1 values were 0.67, 0.63, and 0.64 for BL and 0.45, 0.44, and 0.48 for PB (Supplementary Table S1).

### 3.4. Performance of the common 328K individual-SNP representation

Genomic prediction performance was first evaluated using the common dataset comprising 328,597 individual SNP predictors across 204 rice accessions. Classical machine-learning models, including XGBoost, LightGBM, Random Forest (RF), Support Vector Machine (SVM), were benchmarked against the Quantum Support Vector Classifier (QSVC) for predicting leaf blast (BL) and panicle blast (PB) resistance.

Under binary classification, BL accuracy ranged from 55.5% to 86.5% across the reported SNP-based models, with macro-F1 scores ranging from 0.36 to 0.86. XGBoost, RF, LightGBM, and SVM shared the highest reported BL performance (accuracy = 86.5%, balanced accuracy = 0.86, and macro-F1 = 0.86). For PB, accuracy ranged from 50.2% to 77.0%, with QSVC reaching the highest reported accuracy (77.0%; macro-F1 = 0.73). Comparative binary classification results are presented in Table 2.

Three-class prediction was additionally evaluated using resistant, intermediate, and susceptible categories. Across the reported SNP-based models, macro-F1 scores ranged from 0.31 to 0.69 for BL and from 0.34 to 0.52 for PB (Supplementary Table S1).

### 3.5. Classical machine learning versus quantum-kernel classification

Direct comparisons between classical machine-learning models and QSVC were conducted using matched validation partitions to assess their relative predictive performance. For binary BL prediction, the best-performing classical model, XGBoost, achieved a macro-F1 score of 0.86, compared with 0.82 for QSVC, corresponding to a difference of +0.04. For PB, the corresponding macro-F1 scores were 0.71 and 0.73, respectively (Table 2).

The relative performance of classical and quantum approaches was further examined across the three-class prediction task (Supplementary Table S1). QSVC achieved lower predictive performance relative to the strongest classical baseline, with differences of 0.02 and 0.07 in macro-F1 for BL and PB, respectively.

These results indicate that, under the evaluated feature-selection, quantum-encoding, and validation conditions, quantum-kernel classification the predictive performance of established classical machine-learning approaches. The comparison provides an empirical benchmark for the applicability of QSVC to high-dimensional rice genomic prediction.

### 3.6. Haplotype-based genomic prediction of leaf and panicle blast

The predictive performance of phased and unphased haplotype representations was evaluated using the same disease-response definitions and outer validation partitions applied to the individual-SNP models.

For binary BL prediction, phased-haplotype models achieved accuracy values ranging from 49.1% to 87.3%, with macro-F1 scores ranging from 0.36 to 0.87. The phased SVM had the highest reported BL accuracy (87.3%) and was tied with phased XGBoost for the highest macro-F1 score (0.87). Unphased-haplotype models produced macro-F1 scores ranging from 0.36 to 0.87, with SVM attaining the highest value (0.87).

For PB, the best-performing phased and unphased haplotype models achieved macro-F1 scores of 0.73 and 0.72, respectively. The corresponding balanced accuracy values were 0.75 and 0.75. (Table 2; Figure 3).

Under three-class classification, phased and unphased haplotype models achieved macro-F1 scores of 0.32-0.69 and 0.31-0.68 for BL, respectively. For PB, the corresponding ranges were 0.35-0.53 and 0.37-0.51 (Supplementary Table S1; Figure 3).

Overall, haplotype-based prediction was broadly comparable with individual-SNP prediction, although the direction and magnitude of differences depended on the disease trait, genomic representation, and algorithm.

### 3.7. Direct comparison of SNP and haplotype representations

A direct comparison of individual SNPs, phased haplotypes, and unphased haplotypes was performed to determine the influence of genomic representation on prediction performance.

For BL, the best-performing SNP model achieved a macro-F1 score of 0.86, compared with 0.87 and 0.87 for phased and unphased haplotype representations, respectively. The corresponding values for PB were 0.73, 0.73, and 0.72 (Table 2; Figure 3).

Relative to the SNP representation, phased haplotypes changed the best macro-F1 by +0.01 for BL and 0.00 for PB. Unphased haplotypes produced corresponding changes of +0.01 and -0.01.

Importantly, phased and unphased haplotype representations reduced predictor dimensionality by approximately 68.3% and 85.0%, respectively. Thus, the observed performance differences were considered alongside the substantial reduction in genomic predictor variables.

These results demonstrate that phased haplotypes preserved the best reported binary macro-F1 while using 68.3% fewer predictors than the SNP representation. Unphased haplotypes also reduced predictor dimensionality substantially, but their PB best macro-F1 was 0.01 lower than the SNP and phased-haplotype results. The relative advantages of each representation depended on disease trait and model, highlighting the importance of evaluating genomic encoding as an independent component of prediction model development.

### 3.8. Contrasting predictability of leaf and panicle blast

Prediction performance differed between BL and PB across the evaluated genomic representations and machine-learning approaches. Under binary classification, the best reported macro-F1 scores were 0.87 for BL and 0.73 for PB, a difference of 0.14.

Across the SNP, phased-haplotype, and unphased-haplotype representations, BL had higher best binary macro-F1 performance in all 3 of 3 representation-specific comparisons (Figure 3).

Differences between BL and PB were also observed under three-class classification. The results workbook does not include confusion matrices, so intermediate-class recall and misclassification counts cannot be quantified from the available source.

These findings indicate that genomic predictability was not identical between BL and PB. Potential contributions from trait-specific genetic architecture, environmental effects, and disease-response variation are considered in the discussion.

## 4 DISCUSSION

### 4.1. From GWAS-informed GBLUP to machine-learning genomic prediction of rice blast resistance Akohoue et al. (2026) demonstrated that incorporating genome-wide association study

(GWAS)-derived marker weights into genomic best linear unbiased prediction (GBLUP) improved the prediction of leaf blast (BL) and panicle blast (PB) resistance by up to 37%, highlighting the value of biologically informed genomic prediction in rice. Genomic prediction has become an important component of modern crop improvement because genome-wide marker information can support the prediction and selection of superior genotypes (Crossa et al., 2017). More recently, advances in machine learning and high-dimensional genomic analysis have expanded the range of approaches available for genomic prediction in plant breeding (Crossa et al., 2025).

The present study extends this research in a complementary direction by evaluating whether nonlinear machine-learning algorithms and alternative genomic representations can support accurate classification of phenotypic resistance categories. Rather than modifying marker weights within GBLUP, we compared SNP-, phased-haplotype-, and unphased-haplotype-based representations using classical and quantum machine-learning approaches. This comparison provides insights into the effects of genomic representation and model architecture on rice blast resistance classification (Figure 3).

### 4.2. High-density SNPs as a reference for genomic prediction

The 328,597-SNP dataset provided a common genomic reference for comparing SNP- and haplotype-based prediction of rice blast resistance. By evaluating the same phenotypic classification targets and performance metrics across genomic representations, we examined whether alternative predictor encodings could preserve predictive information while reducing dimensionality (Figure 1; Supplementary Table S2). The broadly comparable binary classification performance across representations suggests that substantial predictor compression can retain useful genomic information for resistance prediction (Table 2).

However, these findings do not establish that increasing marker density necessarily improves predictive performance. The performance of genomic prediction depends on multiple factors, including marker information, genetic relationships, trait architecture, population structure, and prediction-model characteristics (Crossa et al., 2017, 2025). The present analysis therefore supports the 328,597-SNP panel as a dense reference for comparing genomic representations rather than demonstrating an intrinsic advantage of marker density. Future evaluations using different SNP densities and identical validation partitions would help determine the optimal balance between marker coverage, computational requirements, and predictive accuracy.

### 4.3. Predictive value and biological rel**evance of haplotype representations**

Haplotype-based representations achieved predictive performance broadly comparable to SNP-based models while substantially reducing predictor dimensionality. For binary BL classification, the highest macro-F1 score reached 0.87 for both phased and unphased haplotypes, matching the best SNP-based performance. For PB, the highest macro-F1 scores were 0.73 for SNPs and phased haplotypes and 0.72 for unphased haplotypes (Table 2). These findings indicate that haplotypes primarily preserved predictive performance rather than consistently outperforming individual SNP markers.

An important distinction between these representations is their genomic construction. The SNP analysis employed 328,597 markers obtained through linkage disequilibrium (LD) pruning of an original resource containing approximately three million variants. In contrast, haplotypes were constructed from the broader variant resource, enabling the representation of local allelic combinations and linkage relationships that may not be fully captured by individual retained SNPs. Phased and unphased haplotype representations reduced predictor dimensionality by 68.3% and 85.0%, respectively, relative to the SNP matrix (Figure 2; Supplementary Table S2).

**Figure 2.**
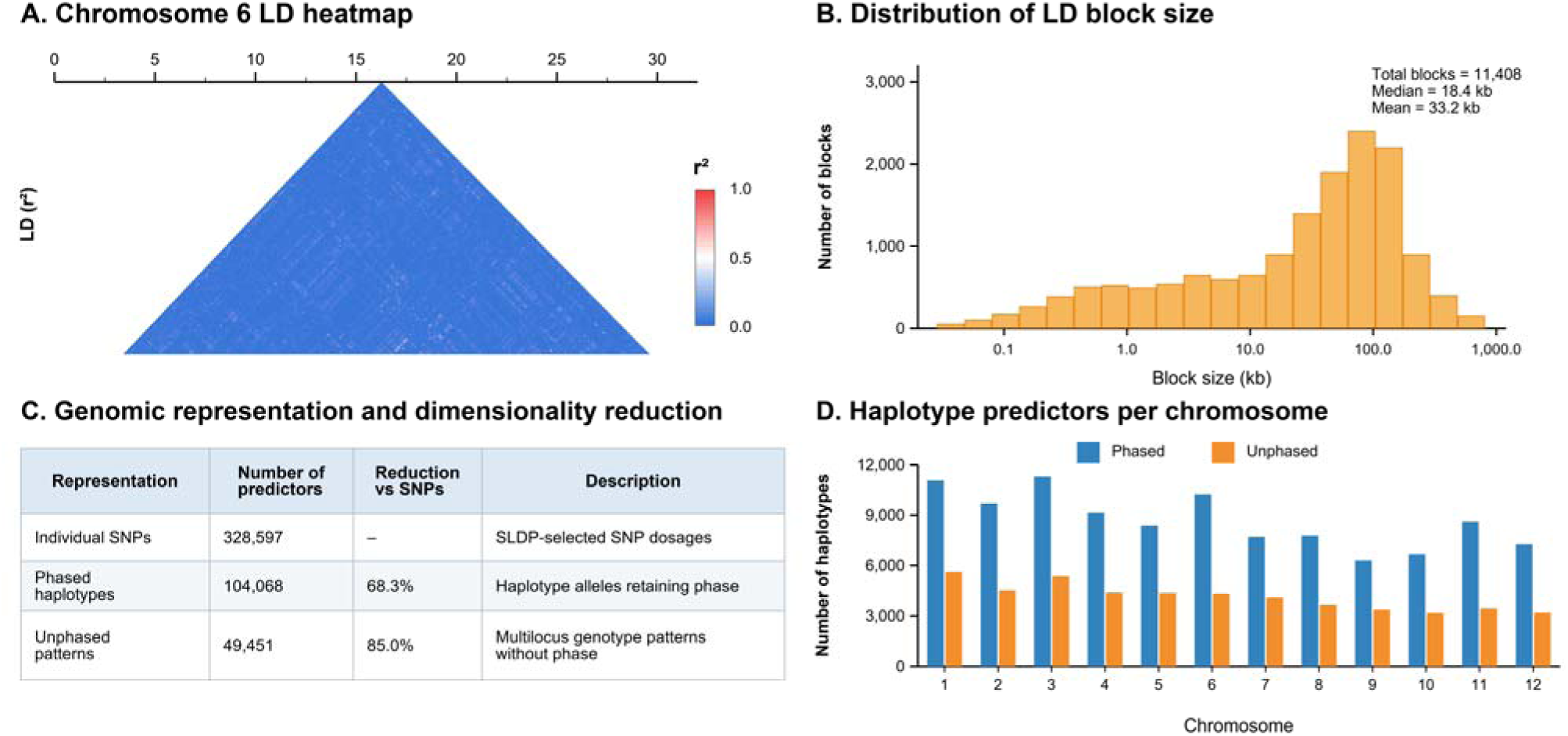
Genome-wide linkage disequilibrium structure, haplotype distribution, and genomic dimensionality reduction. The figure compares individual SNPs with phased and unphased haplotype predictor matrices.

**Figure 3.**
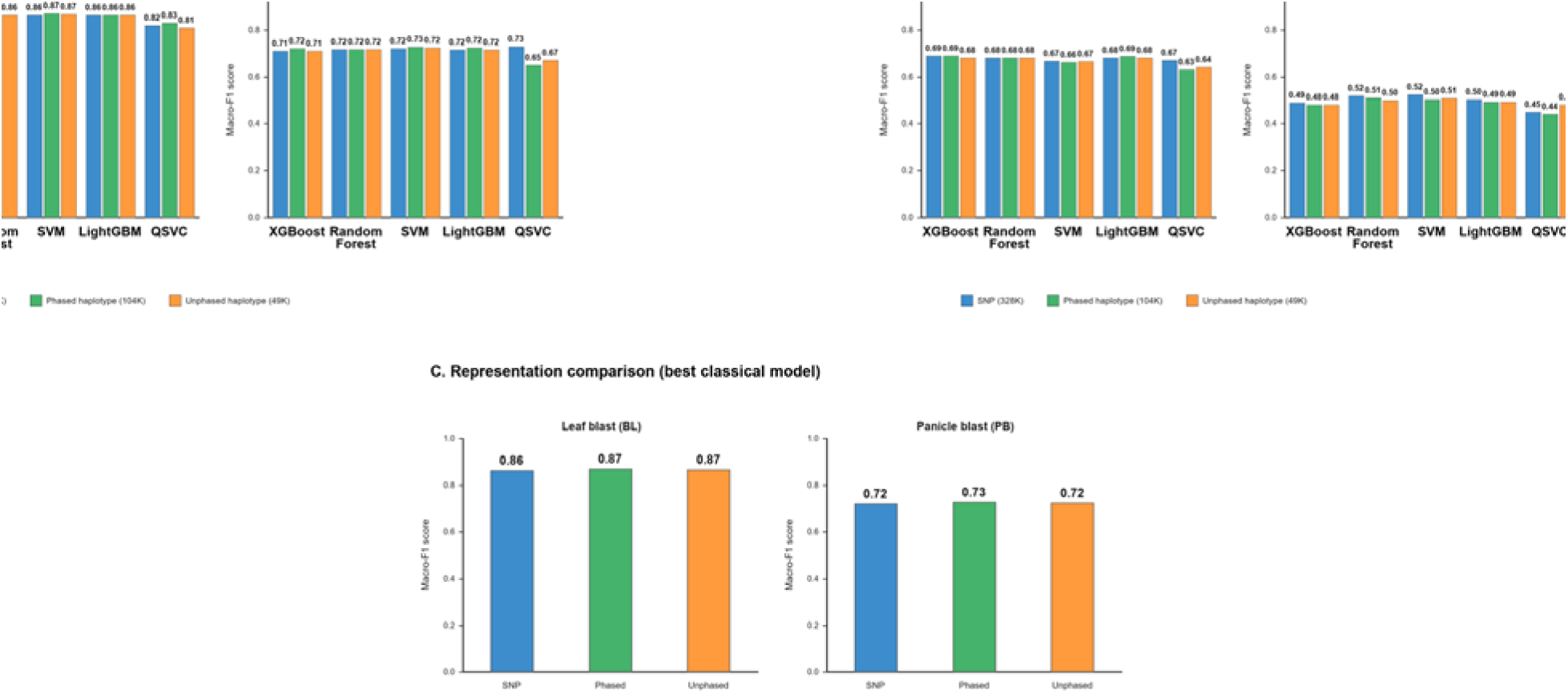
Comparative performance of classical and simulated quantum machine learning for SNP and haplotype-based classification of rice blast resistance. (A) Performance of classical machine-learning models (XGBoost, Random Forest, Support Vector Machine, and Multilayer Perceptron) and simulated quantum support vector classification (QSVC) for binary classification of leaf blast (BL) and panicle blast (PB) resistance using SNP, phased haplotype, and unphased haplotype representations. (B) Comparative performance of the same models and genomic representations for three-class resistance classification (resistant, intermediate, and susceptible). (C) Comparison of the best-performing model pipelines across genomic representations for BL and PB, highlighting differences in predictive performance and predictor dimensionality.

These findings suggest that haplotype-based representations can provide a compact alternative for resistance classification without substantial loss of predictive performance. Their representation of local allelic combinations may also support downstream biological interpretation, integration with transcriptomic and gene-level information, and prioritization of genomic regions for subsequent interaction analyses. Furthermore, reduced predictor dimensionality could be advantageous for computationally intensive applications, including higher-order genetic interaction searches and quantum-inspired optimization. Nevertheless, improvements in computational efficiency or biological interpretability should be evaluated directly rather than inferred solely from reduced predictor counts.

### 4.4. Distinct prediction performance for leaf and panicle blast resistance Although leaf blast (BL) and panicle blast (PB) severity exhibited a significant positive

phenotypic association at the accession level (Spearman’s ρ = 0.65, 95% bootstrap CI = 0.52–0.75; permutation P < 0.00001; Figure 1C), their genomic prediction performance differed substantially. The highest binary macro-F1 score reached 0.87 for BL compared with 0.73 for PB across the evaluated SNP and haplotype representations (Table 2; Figure 3). Three-class classification was more challenging for both traits, particularly PB, which exhibited consistently lower macro-F1 scores (Supplementary Table S1; Figure 3). These findings indicate that a strong phenotypic association between vegetative and reproductive blast responses does not necessarily translate into equivalent genomic predictability.

The contrasting prediction performance may reflect differences in developmental-stage-specific resistance mechanisms, environmental sensitivity, phenotypic reproducibility, and resistance-class distributions. Although BL and PB may share resistance-associated genetic components, their responses are not necessarily governed by identical genetic and physiological processes, consistent with the partially shared but distinct genetic and phenotypic basis of blast resistance described by Akohoue et al. (2026). Nevertheless, the observed phenotypic correlation does not establish identical genetic control, and differences in classification performance alone cannot demonstrate distinct genetic architectures or differences in genetic complexity. Furthermore, unequal resistance-class distributions and variation in classification errors can influence model evaluation, emphasizing the importance of interpreting overall accuracy alongside balanced accuracy and macro-F1 scores (Chicco & Jurman, 2020; Figure 1A–E).

The comparatively lower prediction performance for PB highlights the importance of trait-specific genomic prediction models and continued field-based evaluation of disease responses at both vegetative and reproductive stages. The reduced performance of three-class classification, particularly for PB, also underscores the challenge of predicting resistance at finer phenotypic resolution (Supplementary Table S1). Together, these findings suggest that integrating genomic prediction with complementary phenotypic assessments could improve the reliability of resistance screening and support the development of more robust breeding strategies for durable rice blast resistance.

### 4.5. Classical versus quantum machine learning for genomic prediction

The comparison of classical and quantum machine-learning approaches provides an initial assessment of the feasibility of quantum kernel-based classification for rice blast resistance. Quantum kernel methods use quantum feature representations to construct similarity measures for classification, providing a theoretical basis for quantum-enhanced feature-space learning (Havlíček et al., 2019; Schuld & Killoran, 2019).

In the present study, quantum support vector classification (QSVC) achieved competitive performance in selected binary classification tasks but did not consistently outperform the strongest classical models across traits and genomic representations (Table 2). An important consideration is the difference in feature-processing pipelines. QSVC employed mutual-information-based selection of 64 predictors followed by principal component analysis (PCA) to obtain four components for encoding through a four-qubit quantum feature map (Section 2.7.3). In contrast, the strongest classical models were evaluated using substantially larger representation-specific predictor sets.

Consequently, differences in predictive performance cannot be attributed exclusively to the learning algorithms because the models were not evaluated using equivalent feature information.

The extensive dimensionality reduction required for the quantum implementation may have excluded genomic information relevant to resistance classification. Furthermore, the performance and practical scalability of quantum machine-learning approaches depend on feature encoding, model architecture, trainability, computational resources, and the limitations of available quantum devices (Cerezo et al., 2022).

Finally, the quantum experiments were conducted using simulated circuits rather than quantum hardware (Section 2.7.5). The reported findings therefore establish a pipeline-specific feasibility benchmark rather than evidence of quantum computational advantage. Future investigations should compare classical and quantum models using matched input representations, competitive hyperparameter optimization, and rigorous computational benchmarks before evaluating potential quantum benefits.

### 4.6. Implications for rice blast resistance breeding

SNP- and haplotype-based genomic prediction could complement existing high-throughput phenotyping approaches for rice blast resistance assessment. While UAV-based image analysis is being used to classify disease responses into binary resistance categories, the present study specifically evaluates the prediction of phenotypic resistance classes from genomic markers using classical and quantum machine-learning models.

Genomic prediction has considerable potential to support crop improvement by informing the selection of promising genotypes and improving the efficiency of breeding decisions (Crossa et al., 2017, 2025). In the present study, stronger binary BL classification suggests potential applications in early-stage germplasm screening and prioritization of resistant accessions, whereas comparatively lower PB performance emphasizes the continued importance of field-based disease assessment (Table 2). Haplotype-based representations may provide additional practical value by reducing the number of genomic predictors while maintaining comparable classification performance (Table 2; Supplementary Table S2).

Future integration of SNP- and haplotype-based predictions with UAV-derived disease classifications, multi-omics information, and the GWAS-informed genomic selection framework reported by Akohoue et al. (2026) could strengthen resistance screening, parental selection, and breeding advancement. Such integration will require independent validation across germplasm, environments, and disease-pressure conditions to establish predictive reliability and transferability.

### 4.7. Limitations

Several limitations should be considered when interpreting the findings. First, the genomic dataset comprised 204 unique rice accessions and 2,120 phenotypic observations obtained from repeated evaluations. Although the dataset captured substantial phenotypic variation in leaf and panicle blast resistance, the relatively limited number of genetically independent accessions may constrain the transferability of predictive models to other germplasm populations and environments. Differences in resistance-class distributions may also influence classification performance, particularly for panicle blast (Figure 1; Supplementary Table S1). Independent validation across additional populations and environments will therefore be important for assessing model robustness and generalizability.

Second, the quantum machine-learning experiments were conducted using simulated quantum circuits following substantial dimensionality reduction of the genomic predictors. QSVC employed a four-qubit representation derived from selected genomic features rather than directly processing the complete SNP or haplotype matrices. Consequently, differences in feature dimensionality and preprocessing limit direct comparisons with classical models. Furthermore, simulation-based results cannot establish computational advantage on quantum hardware (Cerezo et al., 2022).

Finally, the present study focused on genomic prediction of phenotypic resistance classes rather than the identification or functional validation of causal resistance-associated variants or higher-order genetic interactions. Future investigations integrating pangenome information, biologically informed feature selection, and experimental validation could help identify genomic regions and interactions contributing to durable rice blast resistance.

### 4.8 Conclusions and Future Directions

This study demonstrates the potential of SNP- and haplotype-based genomic prediction for rice blast resistance classification using a high-density genomic resource. Despite a strong phenotypic association between BL and PB, their contrasting genomic prediction performance highlights the importance of developmental-stage-specific resistance assessment and trait-specific prediction strategies. Binary macro-F1 scores reached 0.87 for BL and 0.73 for PB, while three-class PB classification remained more challenging (Table 2; Supplementary Table S1). Phased and unphased haplotypes reduced predictor dimensionality by 68.3% and 85.0%, respectively, while maintaining broadly comparable predictive performance (Supplementary Table S2). QSVC achieved binary macro-F1 scores of up to 0.83 for BL and 0.73 for PB, demonstrating competitive performance in selected tasks, although classical models generally provided stronger and more consistent benchmarks (Table 2).

Future research could extend these analyses toward pangenome-informed genomic prediction by integrating CT21, Nipponbare, and diverse rice reference genomes to capture structural variants, presence–absence variation, copy-number variation, and divergent resistance-associated haplotypes. Biologically informed screening combined with quadratic unconstrained binary optimization (QUBO), the quantum approximate optimization algorithm (QAOA), and quantum annealing could also be investigated for prioritizing higher-order, three-to five-locus genetic interactions. These approaches require rigorous comparisons with classical methods and independent biological validation, as the present QSVC results establish neither higher-order epistasis nor quantum computational advantage.

For breeding applications, integrating SNP- and haplotype-based genomic prediction with existing UAV-based binary disease classification, high-throughput phenotyping, and multi-omics approaches could strengthen resistance screening and parental selection. Validation across independent germplasm, environments, and disease-pressure conditions will be essential for developing reliable genomic–phenomic prediction pipelines and advancing durable rice blast resistance breeding.

## SUPPLEMENTAL MATERIAL

Supplemental material includes supplementary tables and figures supporting the three-class model evaluation, haplotype and linkage-disequilibrium summaries, and genomic annotation analyses.

## DECLARATION OF GENERATIVE AI AND AI-ASSISTED TECHNOLOGIES

During the preparation of this work, the authors used OpenAI Codex (GPT-5) for language editing and document-formatting assistance. The authors reviewed and edited the content as needed, verified the reported results, and take full responsibility for the content of the published article.

## DATA AVAILABILITY

The genomic and phenotypic datasets analyzed in this study, including SNP and haplotype representations and rice blast resistance observations, are available from the corresponding author upon reasonable request, subject to applicable institutional data-sharing policies.

## AUTHOR CONTRIBUTIONS

Roberto Guerrero: Conceptualization, methodology, software, formal analysis, data curation, visualization, and writing—original draft. Jason Castaño: Software, formal analysis, machine-learning analysis, and writing—review and editing. Félicien Akohoue and Manuel Narvaez: Phenotyping, genotyping, data curation, and writing—review and editing. Gloria Mosquera: Phenotyping, resources, review and editing. Angela Joseph Fernando: Genotyping, data curation, writing—review and editing. Michael Gomez Selvaraj: Conceptualization, methodology, supervision, project administration, resources, and writing—review and editing. All authors read and approved of the final manuscript.

## ETHICAL APPROVAL

This study used rice germplasm and associated genomic and phenotypic datasets and did not involve human participants, animals, or clinical samples. Plant-material handling and associated research activities were conducted in accordance with applicable institutional requirements and Colombian regulations.

## ACKNOWLEDGMENTS

This research was supported by the GI-PPP initiative, funded by the Gates Foundation through the International Institute of Tropical Agriculture (IITA). The authors gratefully acknowledge the Alliance of Bioversity International and the International Center for Tropical Agriculture (CIAT) for institutional support and access to rice genomic and phenotypic research resources. We extend our appreciation to the research and technical teams involved in rice blast phenotyping, genomic data preparation, and machine-learning analyses. We also thank the SuperQ Quantum Computing team for valuable collaborative discussions on quantum and quantum-inspired approaches to higher-order epistasis discovery and acknowledge the use of the Super platform for computational experiments.

## Abbreviations

AE-w: squared additive-effect weighting
BL: leaf blast;
GBLUP: genomic best linear unbiased prediction;
GWAS: genome-wide association study
LD: linkage disequilibrium;
ML: machine learning;
MLP: multilayer perceptron;
PB: panicle blast;
PCA: principal component analysis;
QML: quantum machine learning
QSVC: quantum support vector classifier
RF: random forest
SNP: single-nucleotide polymorphism
SVM: support vector machine
VQC: variational quantum classifier

