## Supplementary table for "Quantum-assisted genomic prediction of blast resistance in rice (Oryza sativa L.) using SNP and haplotype representations"

**Supplementary Table S1.** Comparative performance of classical and quantum machine-learning models for three-class prediction of leaf blast and panicle blast resistance using SNP and haplotype representations

| **Disease trait** | **Genomic representation** | **Model** | **Accuracy (%)** | **Balanced accuracy** | **Macro-F1** |
| --- | --- | --- | --- | --- | --- |
| Leaf blast (BL) | SNP (328,597) | XGBoost | 69.0 | 69.0 | 69.0 |
|  |  | Random Forest | 67.8 | 68.0 | 68.0 |
|  |  | SVM | 66.4 | 67.0 | 67.0 |
|  |  | MLP | NR | NR | NR |
|  |  | QSVC | 67.0 | 66.0 | 67.0 |
|  | Phased haplotypes (104,068) | XGBoost | 68.2 | 68.0 | 69.0 |
|  |  | Random Forest | 67.8 | 68.0 | 68.0 |
|  |  | SVM | 65.9 | 66.0 | 66.0 |
|  |  | MLP | NR | NR | NR |
|  |  | QSVC | 63.0 | 63.0 | 63.0 |
|  | Unphased haplotypes (49,451) | XGBoost | 68.1 | 68.0 | 68.0 |
|  |  | Random Forest | 67.8 | 68.0 | 68.0 |
|  |  | SVM | 66.2 | 66.0 | 67.0 |
|  |  | MLP | NR | NR | NR |
|  |  | QSVC | 63.0 | 63.0 | 64.0 |
| Panicle blast (PB) | SNP (328,597) | XGBoost | 57.3 | 51.0 | 49.0 |
|  |  | Random Forest | 56.4 | 52.0 | 52.0 |
|  |  | SVM | 55.5 | 52.0 | 52.0 |
|  |  | MLP | NR | NR | NR |
|  |  | QSVC | 58.0 | 49.0 | 45.0 |
|  | Phased haplotypes (104,068) | XGBoost | 57.7 | 51.0 | 48.0 |
|  |  | Random Forest | 55.5 | 51.0 | 51.0 |
|  |  | SVM | 53.7 | 50.0 | 50.0 |
|  |  | MLP | NR | NR | NR |
|  |  | QSVC | 54.0 | 47.0 | 44.0 |
|  | Unphased haplotypes (49,451) | XGBoost | 57.7 | 51.0 | 48.0 |
|  |  | Random Forest | 54.2 | 50.0 | 50.0 |
|  |  | SVM | 53.7 | 51.0 | 51.0 |
|  |  | MLP | NR | NR | NR |
|  |  | QSVC | 57.0 | 50.0 | 48.0 |

Three-class prediction was performed using resistant (R; severity scores 1–3), intermediate (I; scores 4–6), and susceptible (S; scores 7–9) categories. Accuracy, balanced accuracy, and macro-F1 were used to evaluate predictive performance. Values should be reported as mean ± standard deviation across cross-validation folds or repeated runs. SNP, phased haplotype, and unphased haplotype representations contained 328,597, 104,068, and 49,451 predictor variables, respectively. Only models actually evaluated should be retained.

**Supplementary Table S2.** Chromosome-Wise Distribution and Genomic Characteristics of Haplotype Blocks in the Rice Diversity Panel

| Chromosome | Number of LD blocks | Mean block length (kb) | Median block length (kb) | Mean SNPs per block | Phased haplotype predictors | Unphased haplotype predictors |
| --- | --- | --- | --- | --- | --- | --- |
| Chr 1 | — | — | — | — | — | — |
| Chr 2 | — | — | — | — | — | — |
| Chr 3 | — | — | — | — | — | — |
| Chr 4 | — | — | — | — | — | — |
| Chr 5 | — | — | — | — | — | — |
| Chr 6 | — | — | — | — | — | — |
| Chr 7 | — | — | — | — | — | — |
| Chr 8 | — | — | — | — | — | — |
| Chr 9 | — | — | — | — | — | — |
| Chr 10 | — | — | — | — | — | — |
| Chr 11 | — | — | — | — | — | — |
| Chr 12 | — | — | — | — | — | — |
| Total | — | — | — | — | 104,068 | 49,451 |
